# Region-level Epimutation Rates in *Arabidopsis thaliana*

**DOI:** 10.1101/2020.08.18.255919

**Authors:** Johanna Denkena, Frank Johannes, Maria Colomé-Tatché

## Abstract

Failure to maintain DNA methylation patterns during plant development can occasionally give rise to so-called ‘spontaneous epimutations’. These stochastic methylation changes are sometimes heritable across generations and thus accumulate in plant genomes over time. Recent evidence indicates that spontaneous epimutations have a major role in shaping patterns of methylation diversity in plant populations. Using single CG dinucleotides as units of analysis, previous work has shown that the epimutation rate is several orders of magnitude higher than the genetic mutation rate. While these large rate differences have obvious implications for understanding genome-methylome co-evolution, the functional relevance of single CG methylation changes remains questionable. In contrast to single CG, solid experimental evidence has linked methylation gains and losses in larger genomic regions with transcriptional variation and heritable phentoypic effects. Here we show that such region-level changes arise stochastically at about the same rate as those at individual CG sites, are only marginal dependent on region size and cytosine density, but strongly dependent on chromosomal location. We also find consistent evidence that region-level epimutations are not restricted to CG contexts but also frequently occur in non-CG regions at the genome-wide scale. Taken together, our results support the view that many differentially methylated regions (DMRs) in natural populations originate from epimutational events and may not be effectively tagged by proximal SNPs. This possibility reinforces the need for epigenome-wide association studies (EWAS) in plants as away to identify the epigenetic basis of adaptive traits.

## Introduction

Cytosine methylation is an epigenetic modification with important roles in the silencing of transposable elements (TEs), the formation of heterochromatin and the regulation of some genes (Kawakatsu et al., 2016). While cytosine methylation in mammals occurs almost exclusively in CG context (CpG), the methylation of cytosines in plants is also abundant in the CHG and CHH contexts (H = C, T or A) (Law and Jacobsen, 2010). In plants, methylation of cytosines in each of the three contexts is maintained by different well-characterized pathways, which are broadly conserved across taxa, suggesting that proper DNA methylation is subject to strong evolutionary constraints (Law and Jacobsen, 2010). Nonetheless, stochastic losses and gains of methylation can arise in plant genomes independently of genetic mutations (Becker et al., 2011; Schmitz et al., 2011), probably as a byproduct of imperfect maintenance fidelity across cell divisions. Once acquired, these so-called ‘spontaneous epimutations’ are heritable over many generations and have the potential to affect the transcriptional output of nearby genes (Schmitz et al., 2013a). Recent analyses of natural populations of *Arabidospis* and maize have provided strong indications that the accumulation of spontaneous epimutations has a major role in shaping methylation diversity patterns over evolutionary time (Vidalis et al., 2016; Xu et al., 2020), although its role in adaptive processes remains unclear (Johannes and Schmitz, 2019; Seymour and Becker, 2017).

Quantitative insights into the formation and trans-generational inheritance of spontaneous epimutations have come from careful studies of Mutation Accumulation (MA) lines. MA-lines are populations of plants that are derived from a single founder and propagated over multiple generations in a stable environment (Becker et al., 2011; Schmitz et al., 2011; Shaw et al., 2000). Using MA-lines of the model plant *A*. *thaliana*, van der Graaf et al. (2015) estimated the rate at which single cytosines in CG context gain methylation (gain rate *α*) at 2.56 · 10^−4^ per generation per halpoid methylome and the loss of methylation (loss rate *β*) at 6.30 · 10^−4^ per generation per haploid methylome. These rates are about 5 orders of magnitude higher than the genetic mutation rate of 6.95 · 10^−9^ calculated for *A*. *thaliana* by Weng et al. (2019). These large rate differences predict that methylome diversity arises much more rapidly than genomic diversity in natural populations, and that epigenetic variation becomes uncoupled from genetic variation over evolutionary time-scales (van der Graaf et al., 2015). One important practical implication of this is that segregating epimutations and their potential functional consequences, are not effectively tagged by proximal SNPs in genome-wide association studies (Johannes et al., 2008).

The relevance of these insights can be questioned on the grounds that there is currently no evidence that methylation status changes at single CGs have any functional consequences in plants. By contrast, gains and losses of methylation over larger regions have been repeatedly linked to heritable phentoypic variation in a number of plant species (Cubas et al., 1999; Gallusci et al., 2016; Ong-Abdullah et al., 2015). Several studies have reported that such regions-level changes (ranging from 50 bp to 1 kb in length) are rare, but do occur in *A*. *thaliana* MA lines at about the same frequencies as genetic mutations (Becker et al., 2011; Ganguly et al., 2017; Hofmeister et al., 2017; Jiang et al., 2014; Schmitz et al., 2011). Some subsequent publications interpreted this to mean that the epimutation rate for regions is comparable with the genetic mutation rate. However, the number of ‘epimutable’ sites in the genome (i.e. regions containing clusters of cytosines) is far smaller than the number of ‘genetically mutable’ sites (i.e. all nucleotides), which would imply that region-level epimutations rates are actually much higher than the genetic mutation rate. Yet, this hypothesis has never been explored formally.

Here, we provide the first estimates of regions-level epimutation rates in the model plant *A*. *thaliana*. Our results show that the rate and spectrum of region-level epimutations is similar to that observed for single CG epimutations. To gain insights into the predictors of spontaneous epimutations, we study the relationship between genome annotation (like genes or transposable elements (TEs)) and epimutation rates. We also investigate epimutation rate levels in distinct chromosomal regions ((peri-)centromeres and chromosome arms) as well as regions of different size and density. Finally, we compare the found epimutation rates to mutation rates in the same genomic regions. Our results have major implications for understanding how linkage disequilibrium (LD) between genetic variants and differentially methylated regions (DMRs) evolves over time, and reinforces the need to carry out methylation-based epigenome-wide association studies (EWAS) in plants.

## Materials and Methods

### Mutation Accumulation lines

To estimate epimutation rates, we used Whole-Genome-Bisulfite-Sequencing (WGBS) data from four previously published *A*. *thaliana* mutation accumulation (MA) data sets (Figure 1). The pedigrees MA1_1, MA1_2 and MA1_3 were generated from the same common Col-0 plant founder MA1, while the pedigree MA2_3 was generated from a different Col-0 founder. All lines were propagated by single seed descent in the green house (Shaw et al., 2000), and one sibling plant was used for propagating the line while another sibling plant was used for sequencing. MA1_1 has been described in Becker et al. (2011). It consists of 12 branches, from which individuals had been sequenced in generations three, 31 and 32 (Figure 1). MA1_2 has been published by Schmitz et al. (2011) and consists of a pedigree with seven branches with individuals measured in generations three and 31 (Figure 1). MA1_3 and MA2_3 were previously published in van der Graaf et al. (2015). MA1_3 consists of only one branch sequenced at nine different generations (Figure 1), while MA2_3 has two branches sequenced at five different generations (Figure 1).

**Figure 1:**
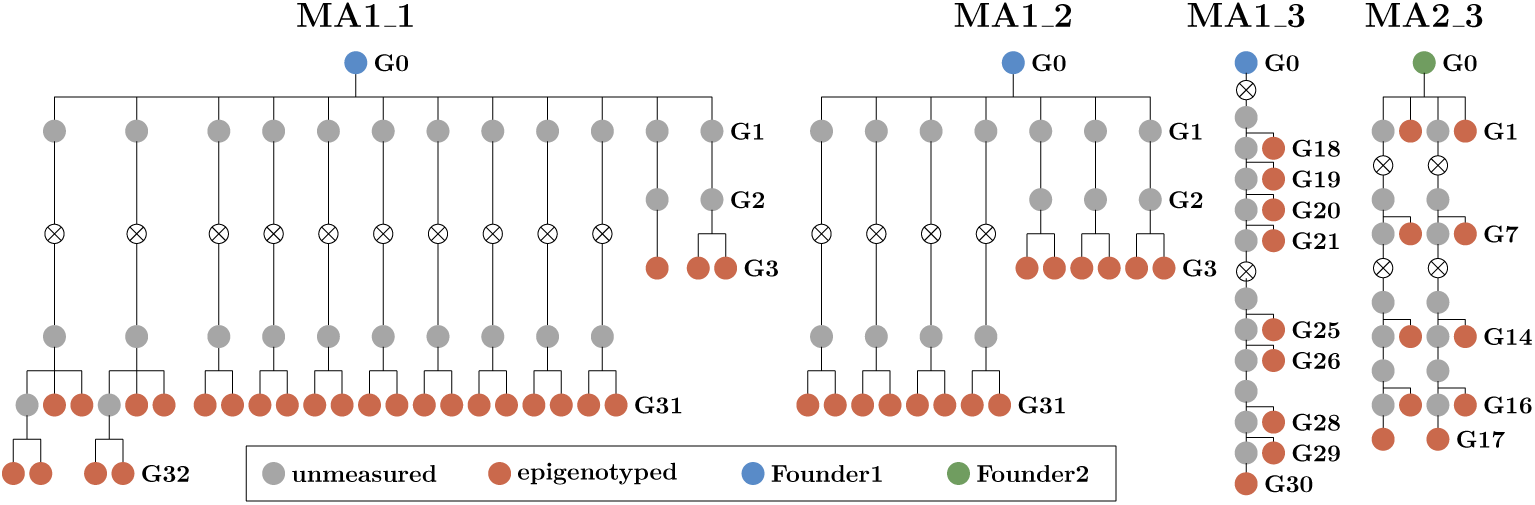
Pedigrees for the Mutation Accumulation lines MA1_1, MA1_2, MA1_3 and MA2_3.

### Constructing Regions

In order to estimate region-level epimutation1 rates, we first defined regions or ‘clusters of cytosines’ based on genomic sequence information. Inspired by approaches used in the animal field for identifying CpG islands, our method starts by defining every pair of cytosines that is a minimal genomic distance apart as seed regions (Figure 2A). In the case of CG, the minimal distance is 0 as every cytosine on the forward strand is complemented by a CG on the backward strand, and there is no nucleotide between the two CGs. For CHG, the minimal distance is 1 and for CHH, because of the lack of symmetry, a seed is formed by every individual CHH on each strand. Then every two seeds with a distance of [minimal distance + 1] between each other are merged. This process is repeated iteratively with increasing distances until either (i) the nearest neighbouring region is more than 185 bp apart, or (ii) the combined region length is higher than 185 bp (Figure 2B and C).

**Figure 2:**
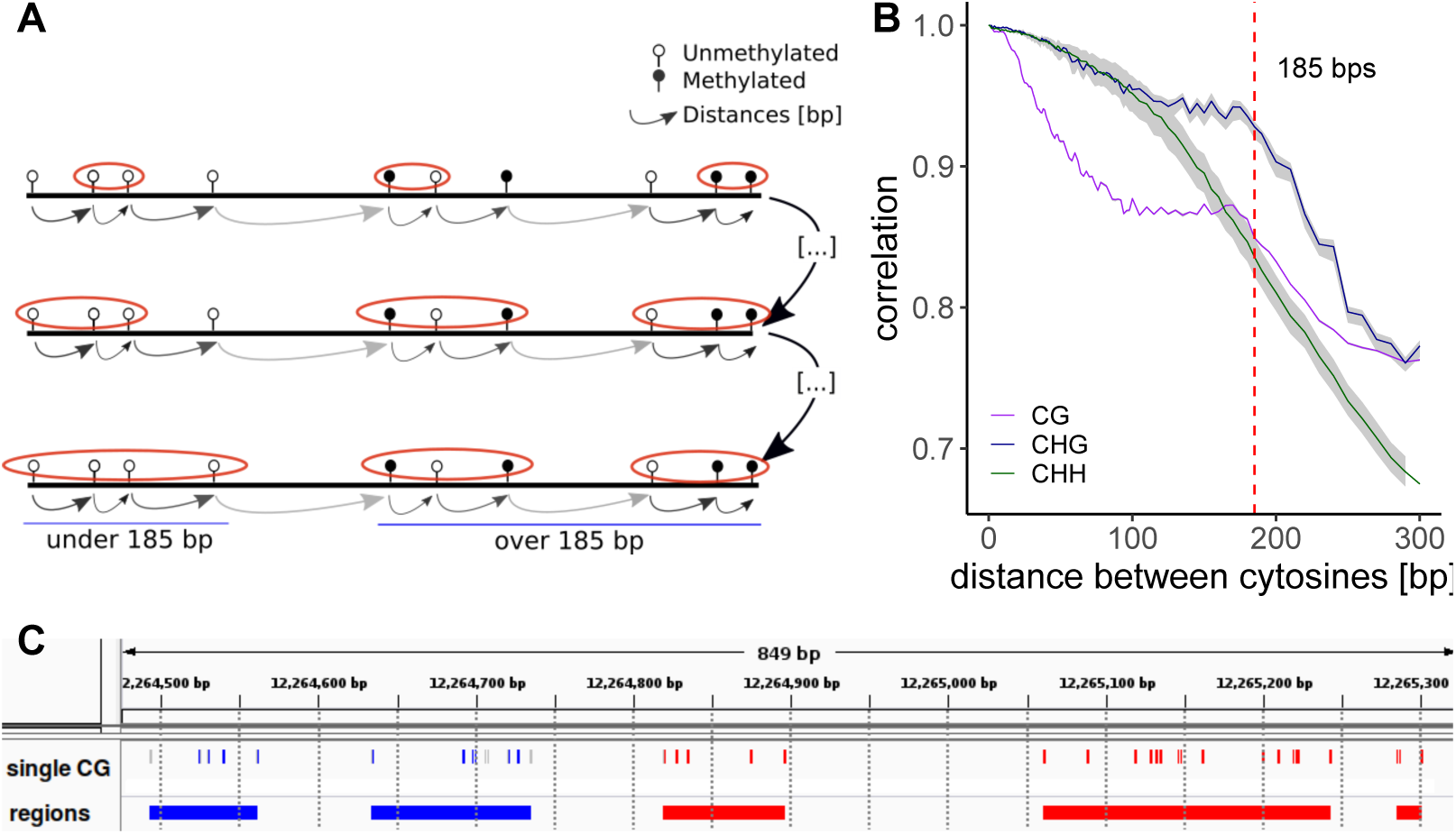
Construction of regions. (**A**) Regions are constructed by iteratively concatenating cytosines (and later clusters of cytosines), if they are less than 185 bp apart and their combined regions size does not exceed 185 bps. The process starts with the cytosines that are closest together and iterates through increasing distances. (**B**) Autocorrelation of cytosines per context for MA line MA1_3. For the autocorrelation of the other MA lines see Figure S1. The grey shading represents the variation caused by the different samples in the MA line pedigrees. (**C**) Example snapshot from the Integrative Genome Viewer(Robinson et al., 2011) showing a stretch of chromosome 1 from MA1_3. The top track shows the single CGs and the bottom track shows the regions. For both tracks, regions/cytosines are either Methylated (blue), Unmethylated (red) or insufficiently covered (gray). For comparison, 100 bp regions are marked with vertical lines, showing how partitioning the genome with an arbitrary window size doesn’t necessarily group close cytosines together.

The 185bp cutoff was based on the decay of the methylation autocorrelation in the genome. We calculated the methylation autocorrelation between neighbouring cytosines per context (CG, CHG or CHH). As expected, the autocorrelation was higher for cytosines at shorter distances compared to cytosines located further away from each other. For contexts CG and CHG the autocorrelation slowly decreased until 185 bp, followed by an abrupt decline (Figure 2B), probably due to the nucleosomic organization of the genome. Therefore we chose 185bp as the cutoff for the constructed regions. For comparison, we also created regions by binning the genome at 100bp, as it is commonly done by DMR calling software (Figure 2C).

### Methylation divergence

After having set up the regions on the basis of genomic information, methylated and unmethylated read counts were summed up per region. To circumvent using regions with poor coverage, only regions with on average more than three reads per cytosine in all individuals of a pedigree were used for further analysis. We called methylation status in regions using the package METHimpute, which provides a method for imputation of WGBS data using a Hidden Markov Model (HMM) (Taudt et al., 2018), taking the number of methylated and unmethyated reads as input. Utilizing its function `callMethylation`, each region was assigned to one of the three methylation status calls: Unmethylated (U), Methylated (M) or Intermediate (I). Methylation status calls were measured by the maximum posterior probability of the HMM. To avoid methylation calls of poor quality only regions with a maximum posterior probability of at least 0.99 in all individuals of a pedigree were used for further analysis. For every pair of individuals in a pedigree we calculated the methylation divergence, with respect to the generation time Δ*t*, which is counted as the independent number of selfing events from their most recent common founder (Figure S2). Following van der Graaf et al. (2015), methylation divergence between two plants *i* and *j, d*_*ij*_, was calculated assuming that for region *n* two individual plants have a divergence of 1 if the tuple of their methylation states, (*m*_*i,n*_, *m*_*j,n*_), is discordant ((M,U) or (U,M)), of ½ if their methylation states are intermediate in one of the two individuals ((M,I), (I,M), (U,I) or (I,U)), and of 0 if the two individuals have the same methylation status ((U,U), (I,I), (M,M)). The genome-wide divergence between two individuals *i* and *j* in a pedigree is then defined as:

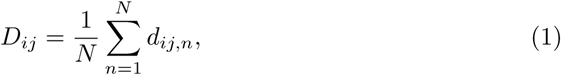

where

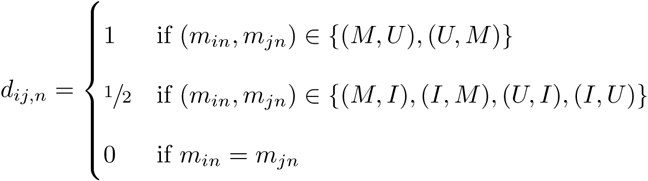

and *N* as the total number of regions.

### Epimutation Rate Estimation

Region-based estimates of the methylation gain rate *α* and the loss rate *β* were obtained with the R-package AlphaBeta Shahryary et al. (2019). We fitted four competing models: ABneutral, ABmm, ABuu, ABnull to the divergence data of each pedigree. Model ABneutral assumes that spontaneous methylation gains and losses accumulate neutrally across generations, ABmm assumes that the accumulation is partly shaped by selection against spontaneous methylation losses, ABuu assumes that the accumulation is partly shaped by selection against spontaneous methylation gains, and ABnull is the null model of no accumulation. Formal model comparisons was carried out as described by Shahryary et al. (2019).

### Annotation specific estimates

We considered methylation accumulation in different genomic regions to investigate which factors had an influence on the epimutation rate estimates. First, regions were categorized by their size and cytosine density. We considered duplets, consisting of two CGs on opposite strands, and longer regions. The longer regions were categorized into high and low cytosine density (above and below the median density of 0.105). Next, we split the regions based on their overlap (≥40% overlap) with genes, transposable elements, 5’-UTRs and 3’-UTRs from The Arabidopsis Information Resource (2018), release TAIR10. Promoters were defined as 1.5kb upstream of each TSS and the genomic space not covered by any of these annotations was defined as intergenic. Genes were further subdivided into gene body methylated genes (gbM genes: mCG enrichment at the gene body, but depletion at Transcription Start and Termination Sites) and non-gbM genes (Bewick et al., 2016). Because of overlapping annotations, some regions correspondeded to more than one annotation. Finally, we tested differences in epimutation rates between centromeres, pericentromeres and chromosome arms. We used the coordinates from Weng et al. (2019), which were adapted from Ossowski et al. (2010) by converting them from TAIR8 to TAIR10.

## Results

### Segmentation

After partitioning the genome based on C density and distance following the strategy outlined above, we found 430 554 CG regions, 473 588 CHG regions and 804 835 CHH regions in the *A*. *Thaliana* genome (Genomic Coordinates in Table S3, S4 and S5). The average number of cytosines per region were 10 (max 66) for CG and CHG and 30 (max 97) for CHH, with a mean distance between cytosines of 9.7bp, 10.4bp and 2.8bp for CG, CHG and CHH regions (Figure 3A). Accordingly, the mean cytosine density per region (i.e. the ratio of base pairs which are C) for CG, CHG and CHH were 0.21, 0.15 and 0.32, respectively (Figure 3B). These observations indicate that most regions were made up of close cytosines.

**Figure 3:**
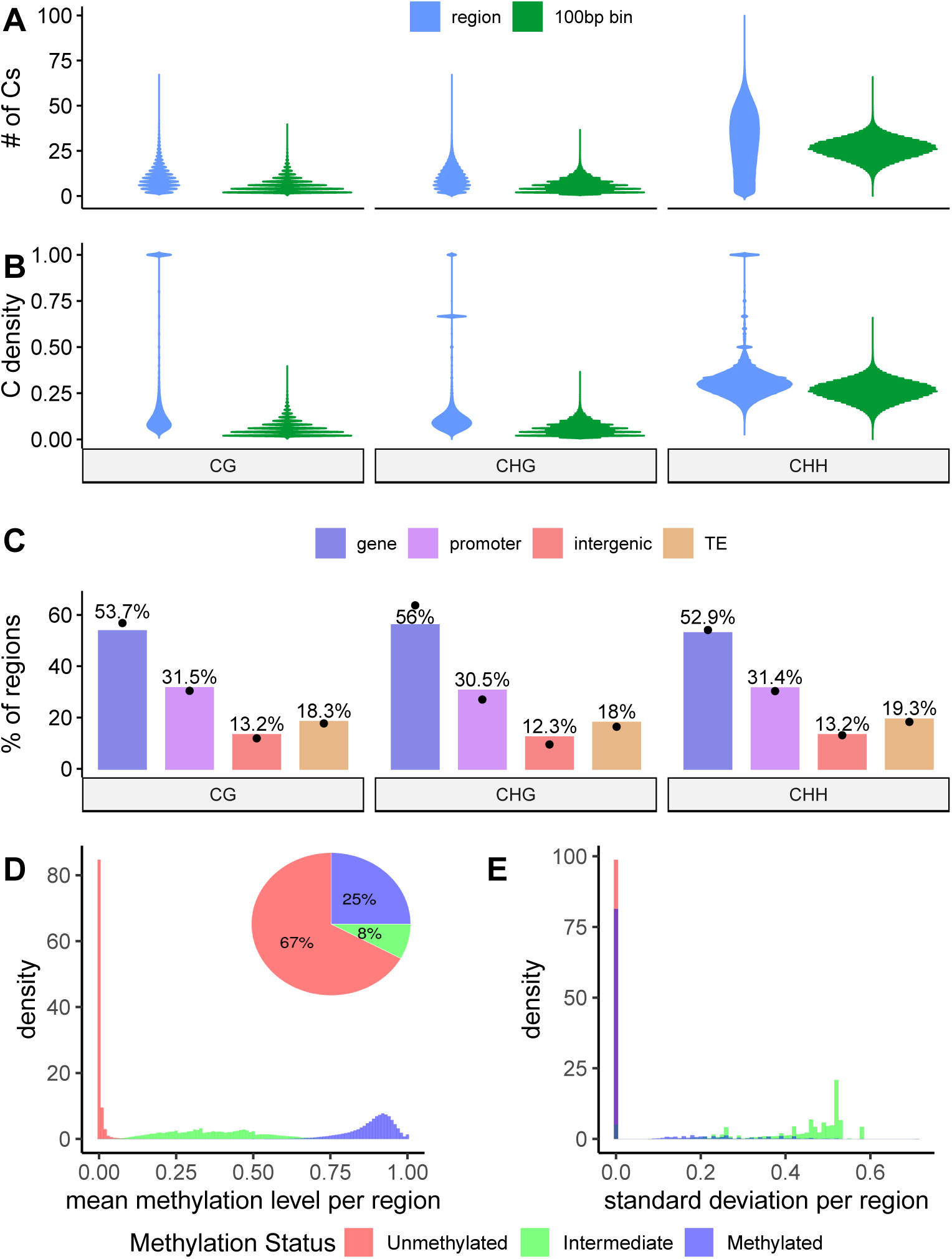
Characteristics of methylation regions. (**A**) Distribution of the number of cytosines per context for regions vs 100bp bins. Mean for regions is 10, 10 and 30 for CG, CHG and CHH. Mean numbers per bins are 5.6, 5.7 and 22.2. (**B**) Distribution of the density of cytosines per context in regions vs. 100bp bins. The mean for regions is 0.21 for CG, 0.15 for CHG and 0.32 for CHH, while for bins it is 0.06, 0.06 and 0.26. (**C**) Percentages of annotations overlapping with regions per context. The dots represent the percentages of single cytosines. (**D**) Mean methylation levels and (**E**) standard deviation per region for all MA lines, colored by whether they were called as Methylated, Unmethylated or Intermediate.

For comparison, we partitioned the genome into 100bp non-overlapping bins. On average, 100bp bins contained less cytosines per bin and consequently lower cytosine densities than the regions (Figure 3A, B). Moreover, 28 % of CG and 21% of CHG bins were made up of only one or two cytosines on either strand, while only 10.1% and 7.1% of CG and CHG regions consisted of a pair of Cs (by construction regions have at least 2 Cs). These higher percentages are relevant, as our aim is to group together close neighboring cytosines instead of constructing a large number of regions with only one or two Cs each. For CHH both the percentage of regions or bins with <= 2 CHHs were low, due to the relative abundance of CHHs throughout the genome (0.008% in bins and 3% in regions).

To study if the genome-wide epimutation rate estimation for regions was not skewed by an over-represented annotation, we calculated the proportion of regions that overlap with a given genomic annotation (Figure 3C). These proportions closely resembled the proportions calculated for single cytosines (dots in Figure 3C). Therefore we would be able to do a meaningful comparison of the epimutation rate estimates between regions and single Cs.

Finally, we explored the methylation levels and status calls per region. First, we studied the extent of the intra-region heterogeneity by calculating a standard deviation over cytosine-level methylation status calls for each region. The I regions showed larger standard deviations compared to U and M regions (Figure 3E and Figure S3B,D), suggesting that U and M regions were very homogeneous, while I regions were a mixture of methylated and unmethylated cytosines. However, the number of I regions were very low in the genome, as on average only ≈ 8%, 4% and 9% of CG, CHG and CHH regions were I. U regions made up the largest proportion of regions in all contexts (≈ 67%, 91% and 89% for CG, CHG and CHH, respectively), while the percentage of methylated regions were highest in CG (≈ 25%), but low in CHG (≈ 6%) and CHH (≈ 2%) (Figure 3D and Figure S3A, C). These results revealed that the constructed regions cluster sets of homogeneous cytosines, that reflect the known genome-wide methylation levels as well as the same proportions of genomic annotations as single cytosines.

### Genome-wide methylation divergence and epimutation rates

We explored whether the regions changed methylation status over time in the same way as single Cs (van der Graaf et al., 2015). We visualized methylation status per region in the more densely sampled pedigrees, MA1_3 and MA2_3 (Figure 1). A number of regions changed methylation status over time and, in general, once a methylation change had taken place it remained stable in subsequent generations (Figure 4).

**Figure 4:**
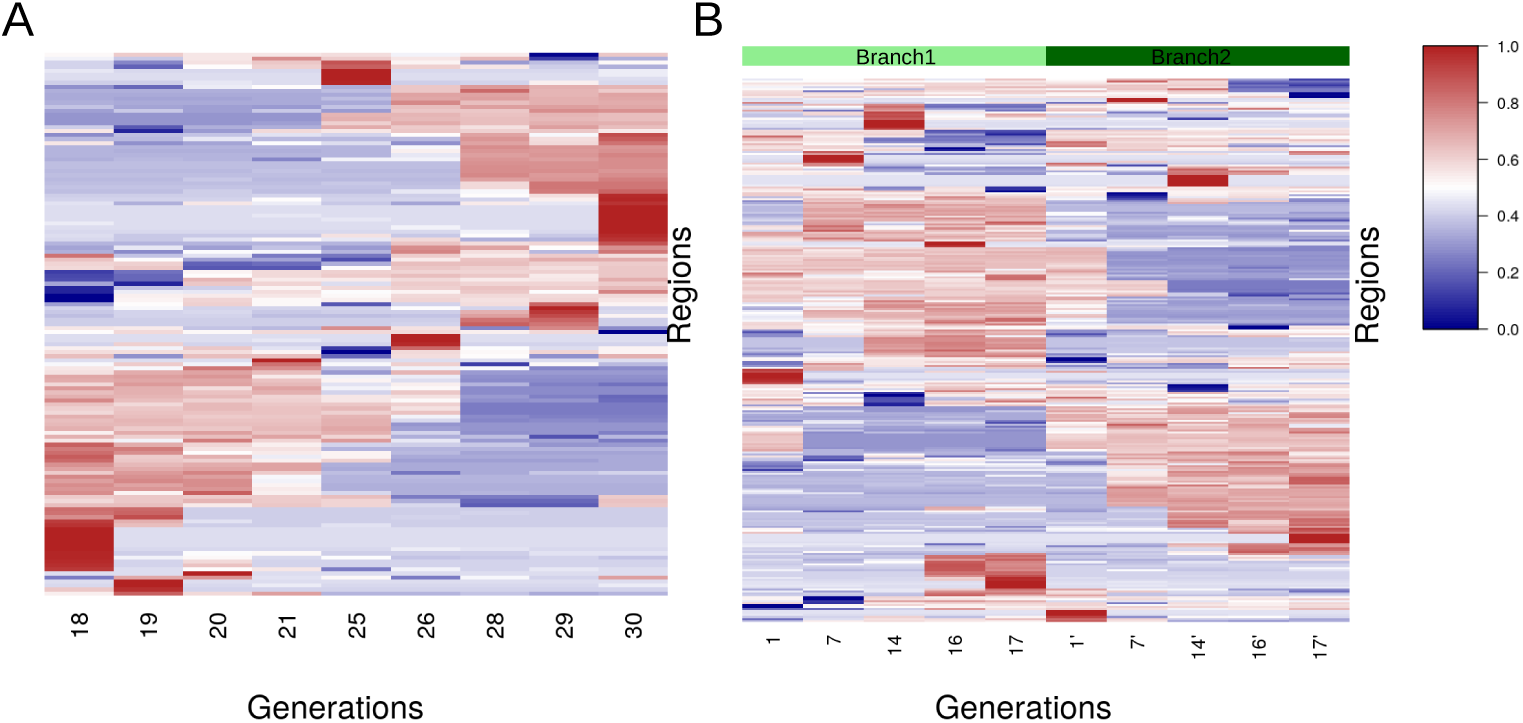
Heatmap of region-wise mean methylation levels over multiple generations of MA line MA1_3 (**A**) and MA2_3 (**B**) for all regions that were assigned both U and M states in at least one individual plant. The regions were clustered using hierarchical clustering.

To quantify these methylation changes genome-wide, we calculated methylation divergence over time using equation 1 and estimated epimutation rates using the model outlined above. We observed that, in all pedigrees, methylation divergence per region in CG context accumulated over time, i.e. two plants which had been selfed separately for longer time (with larger Δ*t*) displayed a higher methylation divergence between them than two plants which were closer to each other in divergence time (Figure 5A). The average CG gain rate *α* was estimated at 1.2·10^−4^ (range: 7.8·10^−5^ - 1.7·10^−4^), while the average CG loss rate *β* was estimated at 4.6·10^−4^ (range: 2.3·10^−4^ - 8.7 · 10^−4^) (Figure 5B).

**Figure 5:**
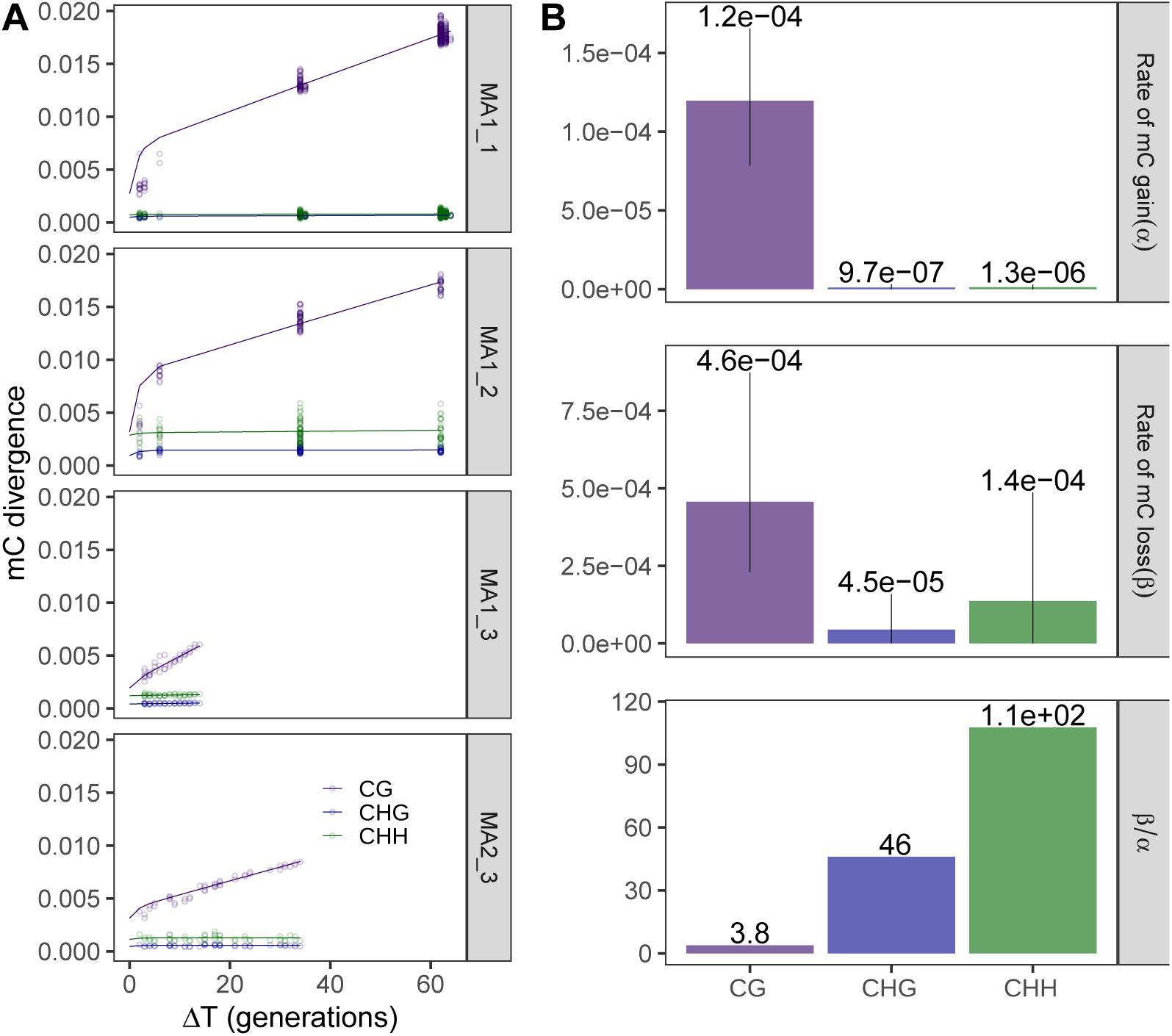
Genome-wide divergence and epimutation rates for CG, CHG and CHH regions. (**A**) mC divergence over generation time for the four MA line pedigrees. (**B**) Mean epimutation rates per context. Error bars represent the maximal and minimal estimates out of the four MA line pedigrees.

Although the divergence profiles of CHG and CHH regions seemed to represent very little accumulation of methylation changes over time, both neutral accumulation model fits were found to be significantly different from the null model of no accumulation (Figure 5A). In accordance with the much lower divergence of CHG and CHH, the epimutation rates were smaller than for CG (for CHG *α* = 9.6 · 10^−7^ (range: 5.0 · 10^−8^ - 2.9 · 10^−6^), *β* = 4.4 · 10^−5^ (range: 6.0 · 10^−7^ - 1.6 · 10^−4^); for CHH *α* = 1.3 · 10^−6^ (range: 4.2 · 10^−9^ - 2.0 · 10^−6^), *β* = 1.4 · 10^−4^ (range: 7.4 · 10^−7^ - 4.9 · 10^−4^), Figure 5B). The higher *β/α* ratios in CHG (46) and CHH (112) in comparison to CG (3.8) reflect the lower genome-wide proportion of methylated regions (see above, Figure S3A and C). The low observed epimutation rates translated into only 33 and 87 regions undergoing a transition from *U* ↔ *M* (on average in the four MA lines, for CHG and CHH respectively), compared to an average of 687.5 CG regions which changed methylation state in the same populations. This low accumulation lead to epimutation rates which were very variable across MA lines. We therefore abstained in the following from splitting these low numbers of regions even further to study annotation-wise epimutation rates in the CHG and CHH contexts.

We tested if a methylation divergence model that considers selection for U or M methylation states fits the data better than the neutral model of epiallele inheritance. Model comparisons revealed that ABneutral provided the best fit to the CG as well as to the CHG and CHH data, indicating that the gain and loss dynamics were neutral in all contexts, at least globally (Table S2). The only exception were contexts CHG and CHH in MA line MA2_3 where models ABuu and ABmm were slightly favored (Table S2); however, this may be an artifact of few and noisy data points.

### Estimation of feature-specific epimutation rates

In addition to genome-wide CG methylation divergence, we also investigated different factors that might have an effect on the rate of methylation change accumulation, namely (1) characteristics of the constructed regions, (2) different genomic annotations and (3) chromosomal organization. To study whether the characteristics of the constructed regions had an effect on methylation loss and gain dynamics, we estimated the epimutation rates for regions of differing size and density. The regions were divided into duplets (2 CGs on opposite strands) and multiplets (regions with multiple CGs), which were further subdivided into dense and sparse based on their C density, as described above. The methylation divergence (Figure 6A) showed a similar accumulation slope for regions of different characteristics, despite the fact that the different regions had variable intercepts (which is attributable to measurement noise (van der Graaf et al., 2015)). Consequently, the epimutation rates were also very similar across the different region types, indicating that region size and cytosine density had no major affect on epimutation rates. However, we did notice a tendency for duplets to lose more and gain less methylation compared to multiplets, resulting in a higher *β/α* ratio. This preferential methylation in CG-rich regions has also been observed using WGBS (Cokus et al., 2008).

**Figure 6:**
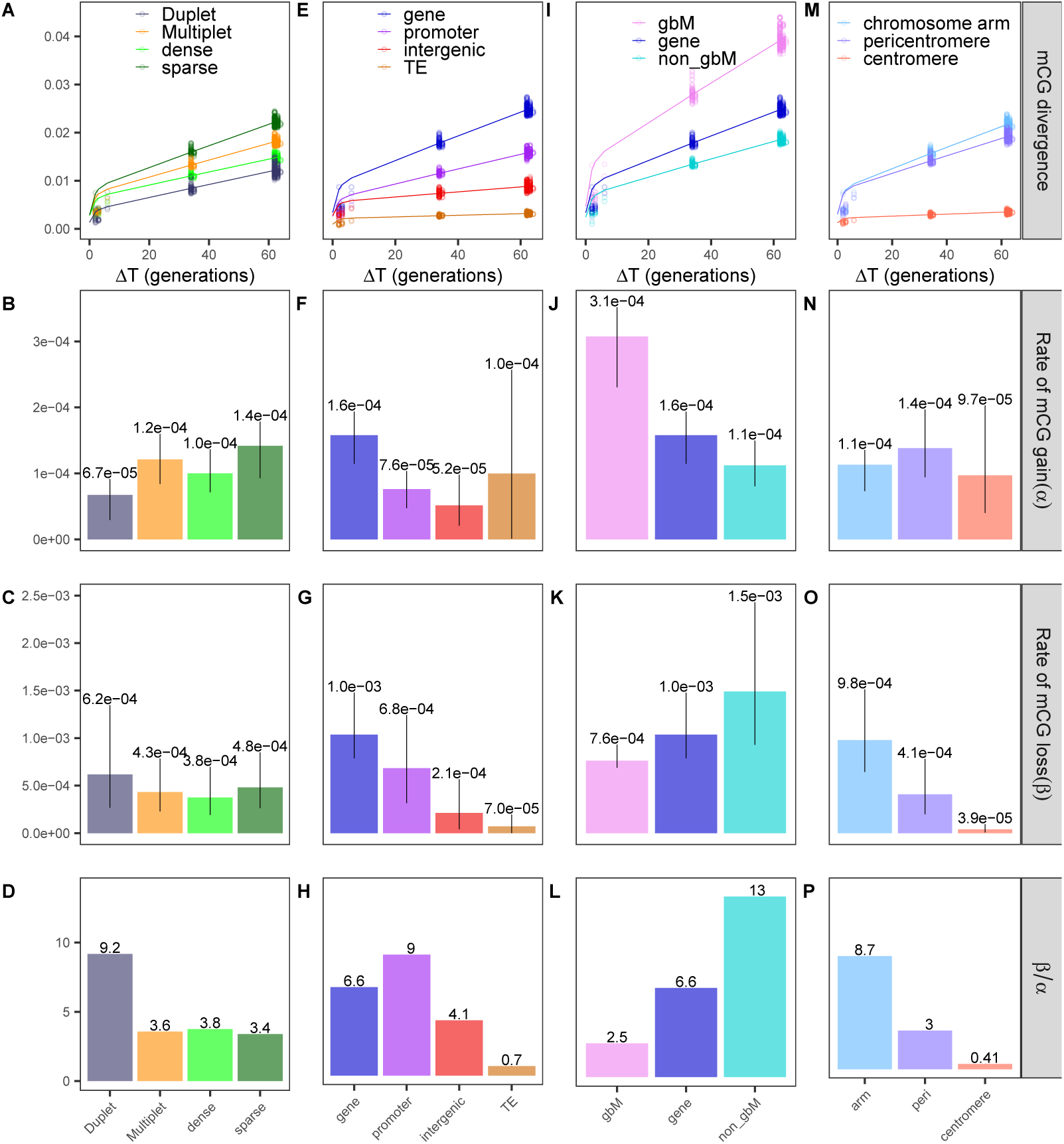
mCG divergence profiles of MA1_1 and mean epimutation rate estimates for different genomic features. Variations of *α* and *β* estimates between MA lines are represented as error bars. All rates per MA line and genomic features are found in table S1. Divergence profiles for all MA lines can be seen in figure S4. (**A**)-(**D**) “Duplet” regions (slategrey) vs. regions with “Multiple” CGs (orange). The regions with multiple CGs were further separated into regions with high (“dense”, bright green) and low density (“sparse”, dark green). (**E**)-(**H**) Genes (blue), promoters (purple), intergenic regions (red) and TEs (darkorange). (**I**)-(**L**) Gene-body methylated genes (violet) vs. non-gbM genes(turquoise). (**M**)-(**P**) Chromosome arms (light blue), pericentromeres (lilac), centromeres (salmon).

Furthermore, we calculated epimutation rates for regions overlapping with genes, promoters, intergenic regions and TEs separately. As observed for single CGs by van der Graaf et al. (2015), the annotation for which regions accumulated the most transgenerational changes was genes, followed by promoters. For intergenic regions and TEs on the other hand the divergence profiles showed very little accumulation (Figure 6E). The average estimated CG epimutation rates are displayed in Figure 6F-H (rates per MA line in Table S1).

Genes exhibited the highest epimutation rates and the ratio *β/α* of 6.6 indicated a strong preference for methylation loss. This could, in part, be attributed to the inability of genic methylation to be recovered once severely compromised (Stroud et al., 2013). Moreover, we split genes into gene body methylated (gbM) genes and non-gbM genes. Interestingly, this more detailed analysis revealed that accumulation of methylation divergence was more rapid in gbM genes than in non-gbM genes (Figure 6I). The epimutation rates of gbM and non-gbM genes resembled a trade-off between methylation gain and loss: while gbM genes gained methylation almost three times as fast as non-gbM genes, they lost methylation at about half the rate of non-gbM genes (Figure 6J, K). The ratio *β/α* of non-gbM genes, which was higher than in any other genomic feature (Figure 6L), implies that while non-gbM genes were kept relatively free of methylation, gbM genes remained considerably more methylated than promoters and intergenic regions, reflecting the higher methylation levels of gbM genes in comparison to non-gbM genes.

Apart from the non-gbM genes, the highest *β/α* ratio was estimated for promoters, where methylation loss was ≈ 9 times higher than methylation gain. As DNA methylation at promoters is often caused by spreading of cytosine methylation from nearby TEs and repeats, gene-adjacent regions like promoters are targeted by the ROS1-dependent active demethylation machinery to prevent transcriptional silencing of genes (Tang et al., 2016; Zhang et al., 2018). TEs, on the other hand, were the only annotation for which gaining methylation was more probable than losing it (*β/α* = 0.7, Figure 6H). This faithfully maintained DNA methylation is a product of targeting by the *de novo* methylation machinery, stemming from the high risk TEs pose to genome integrity (Law and Jacobsen, 2010; Stroud et al., 2013).

Epimutation rates were also calculated for 5’- and 3’-UTRs, although the small number of regions overlapping these annotations made the estimates very variable across MA lines and especially the MA lines with a smaller number of samples (MA1_3 and MA2_3) showed comparatively large standard errors (Table S1). We did observe that the epimutation rates of 3’-UTRs were higher than the rates of 5’-UTRs in all MA lines. Within gene bodies, the 3’ end is the most enriched in CG-DMRs (Schmitz et al., 2011), which could explain the higher epimutation rates in these regions. Both 5’- and 3’-UTRs displayed very high *β/α* ratios (44-87 for 5’- and 10-28 for 3’-UTRs). This tendency towards methylation loss hints at a mechanism that keeps these highly conserved sequences of 5’- and 3-’UTRs, which are critical for controlling mRNA translation and regulating rRNA turnover, respectively, in an unmethylated state (Mignone et al., 2002).

Finally, we explored epimutation rates per chromosome region. mCG divergence was highest for regions in chromosome arms, closely followed by pericentromeres, while centromeres displayed almost no transgenerational changes (Figure 6M). Similar observations were also made by Schmitz et al. (2011), who reported a depletion of CG-DMRs in pericentromeres and centromeres, and by van der Graaf et al. (2015) who for single cytosines found higher mCG divergence in chromosome arms. Estimation of gain and loss epimutation rates showed that all chromosome regions had similar mCG gain rates (Figure 6N), but mCG loss was highest in chromosome arms while lowest in centromeres (Figure 6O). Therefore, while chromosome arms appeared to lose methylation a lot quicker than they gain it, centromeres were the genomic feature that revealed the strongest preference for methylation gain over loss out of all the investigated genomic features (*β/α* = 0.41, Figure 6P). The prevalence of high mCG conservation in centromeres coincides with the distribution of both TEs/repeats and heterochromatin along the genome (Cokus et al., 2008; Schmitz et al., 2013b; van der Graaf et al., 2015), suggesting either confounding with annotation or the influence of heterochromatin-specific methylation factors such as DDM1 (Stroud et al., 2013).

To explore the extent of the confounding between annotations and chromosome regions, epimutation rates were estimated for TEs and genes in chromosome arms, pericentromeres and centromeres separately. From the results it stands out that, in general, a genomic annotation accumulated more epigenetic changes when it was located in chromosome arms compared to centromeres. TEs in chromosome arms even showed a slightly higher mCG loss than gain. In addition, irregardless of chromosome regions, genes were always associated with higher epimutation rates than TEs (Figure 7). Therefore differences in epimutation rates specific to chromosome regions cannot be simply caused by the distribution of TEs vs. genes along the genome. More probably chromatin architecture also plays a relevant role in shaping epimutation rates and epigenomic divergence over time.

**Figure 7:**
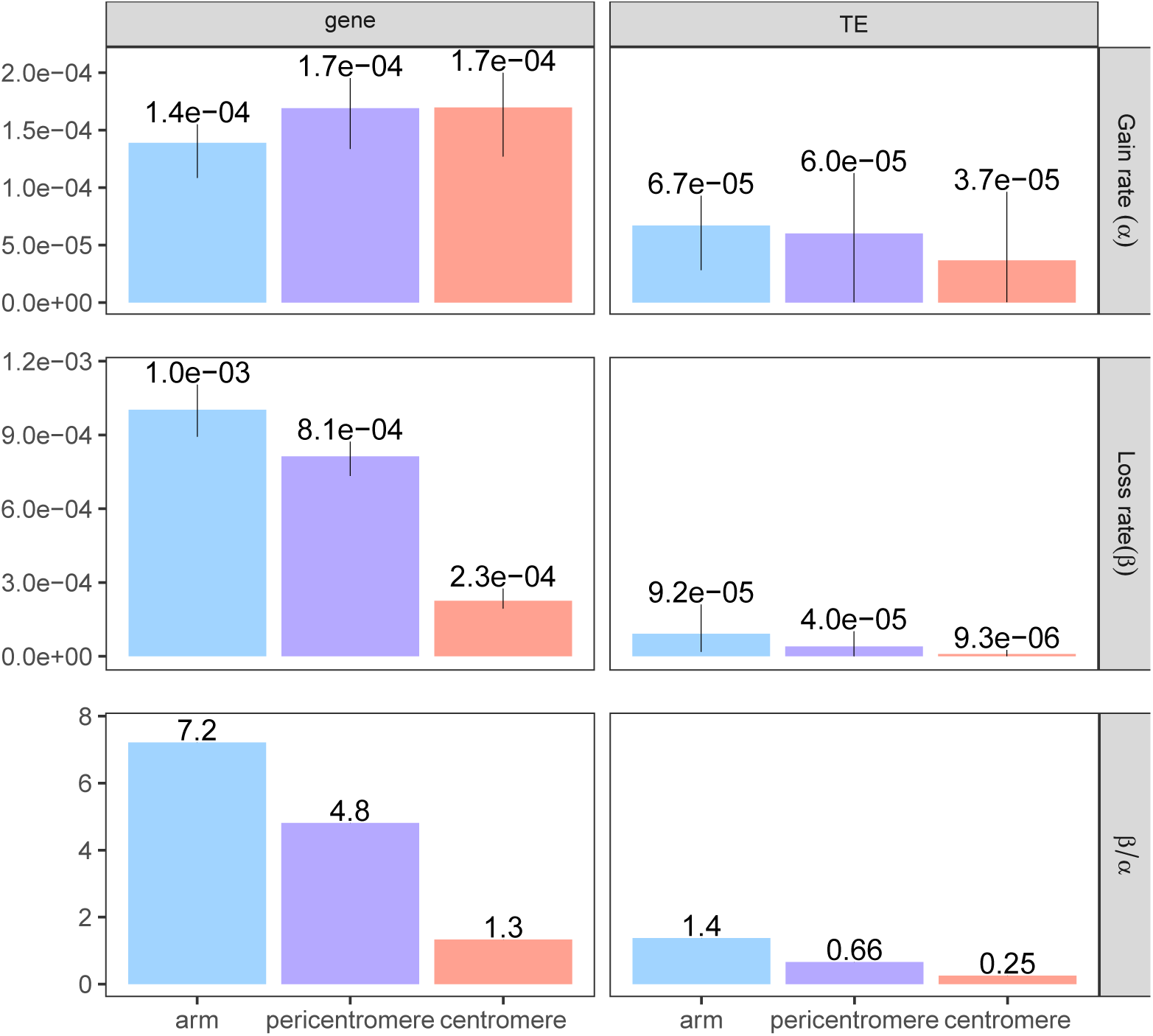
Epimutation rates for TEs and genes in 3 different chromosome regions. Chromosome arms (indigo), pericentromeres (violet), centromeres (salmon). MA line MA1_3 is not shown here as it has a very small number of samples and epimutation rates for this category, with a small number of regions to test, leads to noisy estimates.

As for the genome-wide estimations, we compared the methylation divergence fit for the models with and without selection. In all lines and all annotations, the model without selection could fit the data best (f1-scores and p-values in Table S2). Therefore, if present, genome-wide selection against a methylation state is very low.

## Discussion

In this paper we provide a comprehensive study of region-wise epimutation rates in the model plant *A*. *thaliana*. After construction of methylation regions, we found that epimutation rates for regions were in the same order of magnitude as those observed for single CGs (van der Graaf et al., 2015). This finding indicates that over generation time coordinated changes of methylation involving neighbouring cytosines occur at a comparable rate as individual cytosine changes. These estimations can have far-reaching implications since changes in the methylation status of clusters of cytosines may have more relevant effects on phenotypes than changes in single isolated cytosines, especially when considering regulatory elements such as promoters. Moreover, our model comparison between no selection and selection models showed that, overall, the observed methylation accumulation profiles could be best explained by a model that assumes no genome-wide selection pressure against losing or gaining methylation. Evidence of no selection has been observed in natural populations of *A*. *thaliana* using methylation side frequency spectrum techniques (Vidalis et al., 2016), but only for single cytosines.

We have shown that the rates of region-level mCG changes, while only marginally dependent on region size and cytosine density, strongly depended on genome annotation and are consistently highest in genes and lowest in TEs. This hierarchy of epimutation rates per annotation is not only preserved from single CG estimates in *A*. *thaliana* (van der Graaf et al., 2015), but is also conserved in mitotic studies of epimutation rates in *Populus trichocarpa* (Hofmeister et al., 2019). This suggests that the hierarchy of epimutations is not a product of methylation reinforcement events during seed development but is conserved by DNA methylation maintenance pathways during mitotic cell division upon somatic development in plants (Hofmeister et al., 2019; Johannes and Schmitz, 2019).

The function of CG methylation in genes, especially gene body methylation, still remains elusive, despite hypotheses linking it to transcription (Bewick and Schmitz, 2017). Consistent with the accumulation of gbM over time observed by Bewick and Schmitz (2017), the estimated *α* rates showed that gbM genes gained methylation much more rapidly than non-gbM genes (and any other annotation) whereas non-gbM genes lost methylation quicker (*β*) (Figure 6J). Takuno and Gaut (2013) reported that although classification of gbM genes was conserved among different *A*. *thaliana* MA lines as well as between *O*. *sativa, Z*. *Mays* and *B*. *distachyon*, the methylation of individual CG sites was highly polymorphic, which is consistent with our observed high epimutation rates. The authors proposed a model by which the conservation of gbM across species implies functional relevance and that individual CG sites of methylation may vary without compromising proper gene function as long as the methylation level per gene is kept above a certain threshold. On the other hand, the higher rate of methylation accumulation has been hypothesized to be a passive byproduct of the errant properties inherent to the heterochromatin machinery, rather than a symptom of functional relevancy (Bewick and Schmitz, 2017; Wendte et al., 2019).

Estimation of epimutation rates at chromosomes arms, centromeres and pericentromeres separately showed that epimutation rates were heavily dependent on chromosomal location, with rates that were lowest at centromeres and highest in arms. Although chromosome location is partially confounded with annotation (as arms are gene-rich and centromeres are TE-rich), this could not completely explain the observed trend, as epimutation rates for TEs in chromosome arms were higher than epimutation rates for genes in centromeres (Figure 7). This means that - independently of annotation - specific targeting takes place in centromeres in comparison to chromosome arms. Indeed, TEs in chromosome arms near active genes were shown to be silenced through RdDM (RNA directed DNA METHYLATION), while the silencing of TEs in gene-poor centromeres is dependent on DDM1 (Zemach et al., 2013). Since RdDM requires the nucleosome remodeler DRD1 to maintain DNA methylation and DRD1 remodels heterochromatic nucleosomes less effectively (Zemach et al., 2013), this may lead to less faithfully maintained methylation in TEs located in chromosome arms compared to TEs in centromeres.

Also using *A*. *thaliana* Col-0 MA lines from Shaw et al. (2000), Weng et al. (2019) recently published a genome-wide mutation rate of 6.95 · 10^−9^ per site and generation. The genome-wide epimutation rates for CG regions were 5 orders of magnitude higher than this mutation rate. This finding indicates that, similarly to single Cs, region-level epimutations occur independently of genetic mutations and provide a separate source of genome evolution and diversity (Monroe et al., 2020). Our results have major implications for understanding how linkage disequilibrium (LD) between genetic variants and differentially methylated regions (DMRs) evolves over time, and substantiate the need for methylation-based epigenome-wide association studies (EWAS) in plants.

Weng et al. (2019) also calculated, for the first time, mutation rates for different genomic annotations, distinguishing between TEs, genic and intergenic regions. The mutation rate for TEs (1.36 · 10^−8^) was nearly four times higher than that of genes (3.35 · 10^−9^) and the mutation rate of intergenic regions was almost twice as high as the genic mutation rate (5.75 · 10^−9^). Interestingly, these mutation values displayed the complete opposite trend as observed for epimutations, where changes were lowest in TEs and highest in genes (Figure 8). The same trend was observed when splitting the genome into chromosome arms and (peri-)centromeres: The mutation rates per chromosome region reported by Weng et al. (2019) showed that the mutation rate was more than twice as high in (peri-)centromeric regions as in chromosome arms. These observations reflect large genome-wide differences between genetic and epigenetic divergence, and have implications for understanding genome-methylome co-evolution.

**Figure 8:**
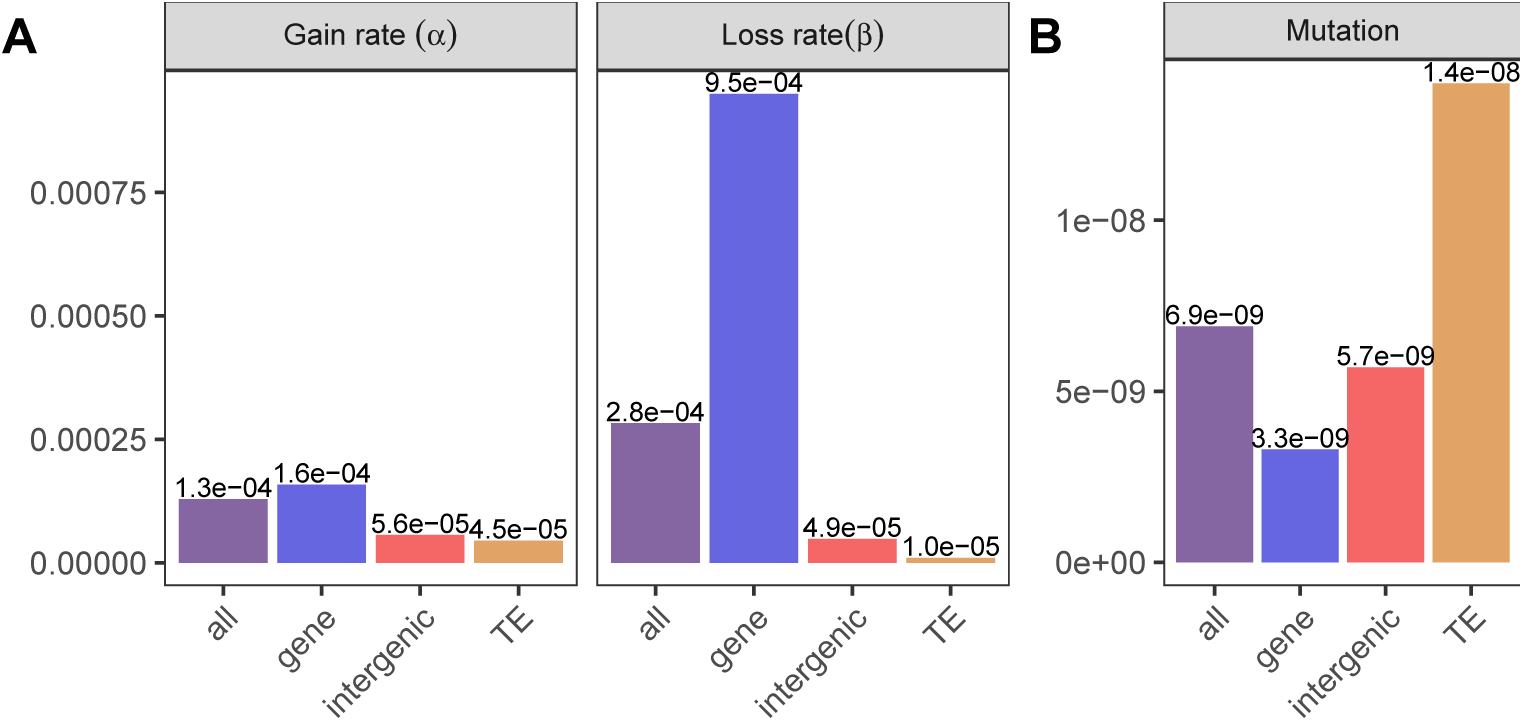
Comparison of region-level epimutation rates from MA1_1 (**A**) with mutation rates calculated by Weng et al. (2019) (**B**).

This study presents the first estimates for region-level epimutation rates for *A*. *thaliana*. The observed methylation changes and subsequent epimutation rates showed that methylation changes of clusters of cytosines accumulate almost as fast as in single cytosines, suggesting that they are independent of genetic change. In addition, we have shown that annotations (genes, promoters, TEs) and chromosome regions are highly predictive of the epimutation rates. Genes and chromosome arms feature the highest accumulation of region methylation changes, while and TEs in centromeres have the lowest epimutation rates. Comparing our epimutation rate estimates to the mutation rates estimated by Weng et al. (2019) we uncovered an anti-correlation between the two, where mutations are lowest where epimutations are highest.

## Acknowledgements

This work was supported by the Impuls-und Vernetzungsfonds of the Helmholtz-Gemeinschaft (grant VH-209 NG-1219).

## Conflict of Interest

All authors declare no conflict of interest.

## Data Archiving

The data, in the form of processed count matrices, was downloaded from GEO with the accession number GSE64463, where the lines MA1_3 and MA2_3 are located in the main GSE matrix and the lines MA1_1 and MA1_2 can be found in the supplementary files. MA1_3 and MA2_3 are described in detail by van der Graaf et al. (2015), while MA1_1 is described in Becker et al. (2011) and MA1_2 in Schmitz et al. (2011).

## Notes

https://www.ncbi.nlm.nih.gov/geo/query/acc.cgi?acc=GSE64463

### Competing Interest Statement

The authors have declared no competing interest.

## References

Becker, C., Hagmann, J., Müller, J., Koenig, D., Stegle, O., Borgwardt, K., et al. (2011). Spontaneous epigenetic variation in the Arabidopsis thaliana methylome. Nature, 480(7376):245–249.

Bewick, A. J., Ji, L., Niederhuth, C. E., Willing, E. M., Hofmeister, B. T., Shi, X., et al. (2016). On the origin and evolutionary consequences of gene body DNA methylation. Proceedings of the National Academy of Sciences of the United States of America, 113(32):9111–9116.

Bewick, A. J. and Schmitz, R. J. (2017). Gene body DNA methylation in plants. Current Opinion in Plant Biology, 36:103–110.

Cokus, S. J., Feng, S., Zhang, X., Chen, Z., Merriman, B., Haudenschild, C. D., et al. (2008). Shotgun bisulphite sequencing of the Arabidopsis genome reveals DNA methylation patterning. Nature, 452(7184):215–219.

Cubas, P., Vincent, C., and Coen, E. (1999). An epigenetic mutation responsible for natural variation in floral symmetry. Nature, 401:157–161.

Gallusci, P., Hodgman, C., Teyssier, E., and Seymour, G. B. (2016). DNA methylation and chromatin regulation during fleshy fruit development and ripening. Frontiers in Plant Science, 7(JUNE2016):1–14.

Ganguly, D. R., Crisp, P. A., Eichten, S. R., and Pogson, B. J. (2017). The arabidopsis DNA methylome is stable under transgenerational drought stress. Plant Physiology, 175(4):1893–1912.

Hofmeister, B. T., Denkena, J., Colomé-Tatché, M., Shahryary, Y., Hazarika, R., Grimwood, J., et al. (2019). The somatic genetic and epigenetic mutation rate in a wild long-lived 2 perennial Populus trichocarpa. bioRxiv, pages 1–56.

Hofmeister, B. T., Lee, K., Rohr, N. A., Hall, D. W., and Schmitz, R. J. (2017). Stable inheritance of DNA methylation allows creation of epigenotype maps and the study of epiallele inheritance patterns in the absence of genetic variation. Genome Biology, 18(1):1–16.

Jiang, C., Mithani, A., Belfield, E. J., Mott, R., Hurst, L. D., and Harberd, N. P. (2014). Environmentally responsive genome-wide accumulation of de novo Arabidopsis thaliana mutations and epimutations. Genome Research, 24(11):1821–1829.

Johannes, F., Colot, V., and Jansen, R. C. (2008). Epigenome dynamics: A quantitative genetics perspective. Nature Reviews Genetics, 9(11):883–890.

Johannes, F. and Schmitz, R. J. (2019). Spontaneous epimutations in plants. New Phytologist, 221(3):1253–1259.

Kawakatsu, T., Huang, S.-s. C., Jupe, F., Sasaki, E., Schmitz, R. J., Urich, M. A., et al. (2016). Epigenomic Diversity in a Global Collection of Arabidopsis thaliana Accessions. Cell, 166:492–505.

Law, J. A. and Jacobsen, S. E. (2010). Establishing, maintaining and modifying DNA methylation patterns in plants and animals. Nature Reviews Genetics, 11(3):204–220.

Mignone, F., Gissi, C., Liuni, S., and Pesole, G. (2002). Untranslated regions of mRNAs. Genome Biology, 3(3):1–10.

Monroe, J. G., Srikant, T., Carbonell-Bejerano, P., Exposito-Alonso, M., Weng, M.-L., Rutter, M. T., et al. (2020). Mutation bias shapes gene evolution in Arabidopsis thaliana. bioRxiv, page 2020.06.17.156752.

Ong-Abdullah, M., Ordway, J. M., Jiang, N., Ooi, S. E., Kok, S. Y., Sarpan, N., et al. (2015). Loss of Karma transposon methylation underlies the mantled somaclonal variant of oil palm. Nature, 525(7570):533–537.

Ossowski, S., Schneeberger, K., Lucas-Lledó, J. I., Warthmann, N., Clark, R. M., Shaw, R. G., et al. (2010). The Rate and Molecular Spectrum of Spontaneous Mutations in Arabidopsis thaliana. Science, 327(5961):1–9.

Robinson, J. T., Thorvaldsdóttir, H., Winckler, W., Guttman, M., Lander, E. S., Getz, G., et al. (2011). Integrative Genome Viewer. Nature Biotechnology, 29(1):24–6.

Schmitz, R. J., He, Y., Valdés-López, O., Khan, S. M., Joshi, T., Urich, M. A., et al. (2013a). Epigenome-wide inheritance of cytosine methylation variants in a recombinant inbred population. Genome Research, 23(10):1663–1674.

Schmitz, R. J., Matthew D. Schultz, Lewsey, M. G., O’Malley, R. C., Urich, M. A., Libiger, O., et al. (2011). Transgenerational Epigenetic Instability Is a Source of Novel Methylation Variants. Science, 334(October):369–373.

Schmitz, R. J., Schultz, M. D., Urich, M. A., Nery, J. R., Pelizzola, M., Libiger, O., et al. (2013b). Patterns of population epigenomic diversity. Nature, 495(7440):193–198.

Seymour, D. K. and Becker, C. (2017). The causes and consequences of DNA methylome variation in plants. Current Opinion in Plant Biology, 36:56–63.

Shahryary, Y., Symeonidi, A., Hazarika, R. R., Denkena, J., Mubeen, T., Hofmeister, B., et al. (2019). AlphaBeta: Computational inference of epimutation rates and spectra from high-throughput DNA methylation data in plants. bioRxiv.

Shaw, R. G., Byers, D. L., and Darmo, E. (2000). Spontaneous Mutational Effects on Reproductive Traits of. Genetics, 155:369–378.

Stroud, H., Greenberg, M., and Feng, S. (2013). Comprehensive analysis of silencing mutants reveals complex regulation of the Arabidopsis methylome. Cell, 152:352–364.

Takuno, S. and Gaut, B. S. (2013). Gene body methylation is conserved between plant orthologs and is of evolutionary consequence. Proceedings of the National Academy of Sciences of the United States of America, 110(5):1797–1802.

Tang, K., Lang, Z., Zhang, H., and Zhu, J. K. (2016). The DNA demethylase ROS1 targets genomic regions with distinct chromatin modifications. Nature Plants, 2(November):1–10.

Taudt, A., Roquis, D., Vidalis, A., Wardenaar, R., Johannes, F., and Colome-Tatché-Tatché, M. (2018). METHimpute: Imputation-guided construction of complete methylomes from WGBS data. BMC Genomics, 19(1):1–14.

The Arabidopsis Information Resource (2018). ftp://ftp.arabidopsis.org/home/tair/Genes/TAIR10_genome_release/TAIR10_gff3/.on www.arabidopsis.org. accessed 30-August-2018.

van der Graaf, A., Wardenaar, R., Neumann, D. A., Taudt, A., Shaw, R. G., Jansen, R. C., et al. (2015). Rate, spectrum, and evolutionary dynamics of spontaneous epimutations. Proceedings of the National Academy of Sciences, 112(21):6676–6681.

Vidalis, A., Zivkovic, D., Wardenaar, R., Roquis, D., Tellier, A., and Johannes, F. (2016). Methylome evolution in plants. Genome Biology, 17:1–14.

Wendte, J. M., Zhang, Y., Ji, L., Shi, X., Hazarika, R. R., Shahryary, Y., et al. (2019). Epimutations are associated with CHROMOMETHYLASE 3-induced de novo DNA methylation. eLife, 8:1–27.

Weng, M.-l., Becker, C., Hildebrandt, J., Neumann, M., Rutter, M. T., Shaw, R. G., et al. (2019). Fine-Grained Analysis of Spontaneous Mutation Spectrum and Frequency in Arabidopsis thaliana. Genetics, 211(February):703–714.

Xu, G., Lyu, J., Li, Q., Liu, H., Wang, D., Zhang, M., et al. (2020). Adaptive evolution of DNA methylation reshaped gene regulation in maize. bioRxiv, pages 1–25.

Zemach, A., Kim, M. Y., Hsieh, P. H., Coleman-Derr, D., Eshed-Williams, L., Thao, K., et al. (2013). The arabidopsis nucleosome remodeler DDM1 allows DNA methyl-transferases to access H1-containing heterochromatin. Cell, 153(1):193–205.

Zhang, H., Lang, Z., and Zhu, J.-k. (2018). Dynamics and function of DNA methylation in plants. Nature Reviews Molecular Cell Biology, 19(August):489–509.

