## Supplementary material for "Region-level Epimutation Rates in *Arabidopsis thaliana*": Figure S

### Region-level Epimutation Rates in *Arabidopsis Thaliana*: Supplementary Information

**Author Details:**

[1] Institute of Computational Biology, Helmholtz Zentrum München Neuherberg 85764, Germany

[2] Department of Plant Sciences, Hans Eisenmann-Zentrum for Agricultural Sciences, Technical University Munich, Freising, Germany

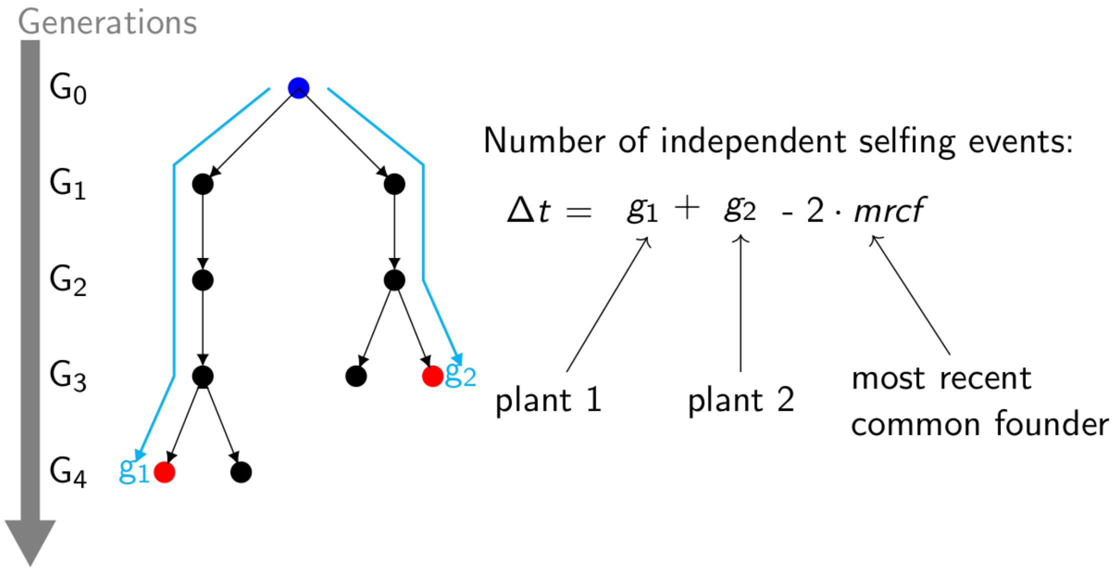


**Figure S1:** Schematic overview on how to calculate $\Delta t$.

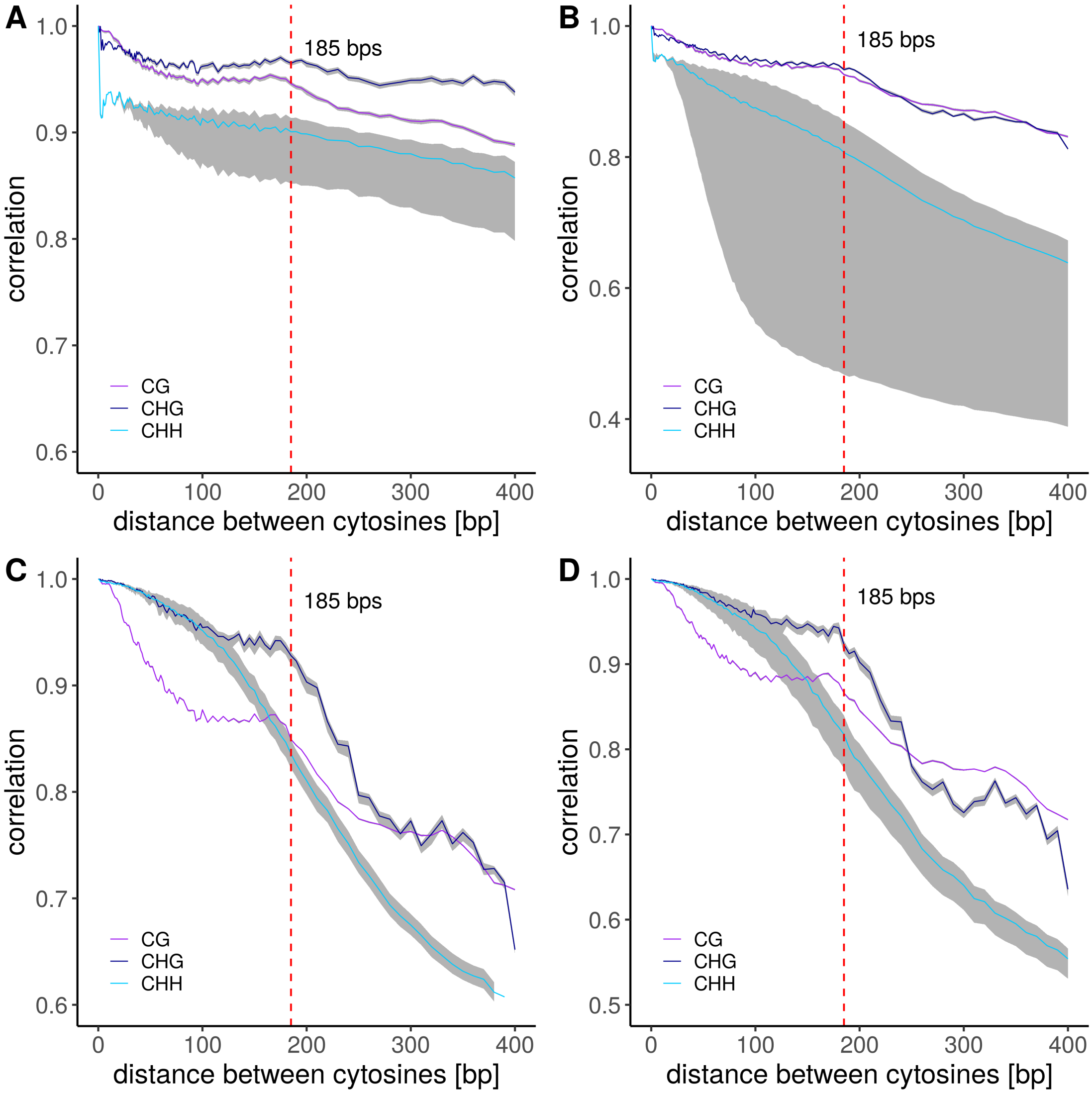


**Figure S2:** Autocorrelation of methylation states per context in MA line (**A**) MA1_1, (**B**) MA1_2, (**C**) MA1_3, (**D**) MA2_3. The grey shading around the lines represents the variation of the samples per MA line.

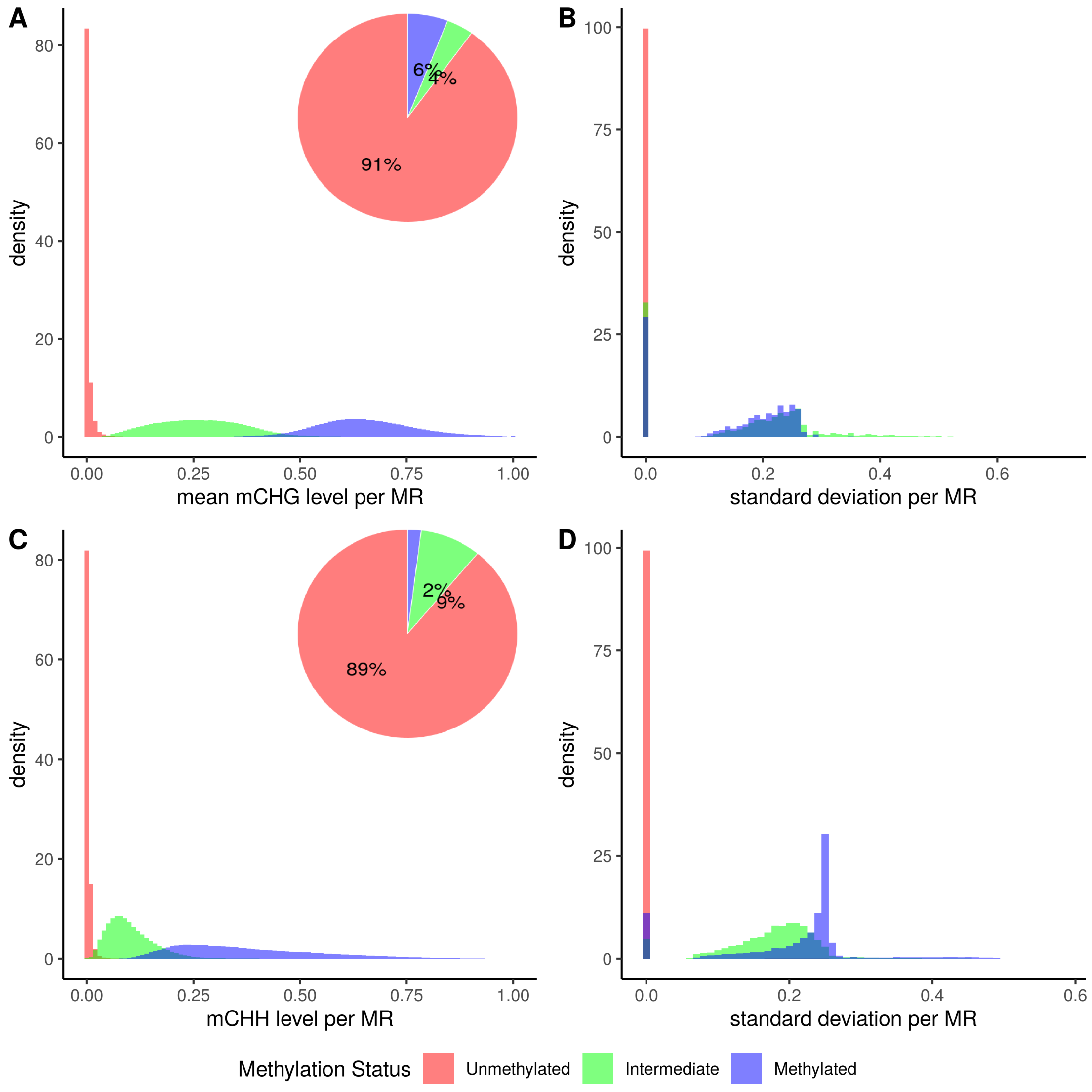


**Figure S3:** **(A)** Mean methylation levels per region for CHG context. **(B)** Standard deviation per region for CHG context. **(C)** Mean methylation levels per region for CHH context. **(D)** Standard deviation per region for CHH context. Colored by whether they were called as Methylated, Unmethylated or Intermediate.

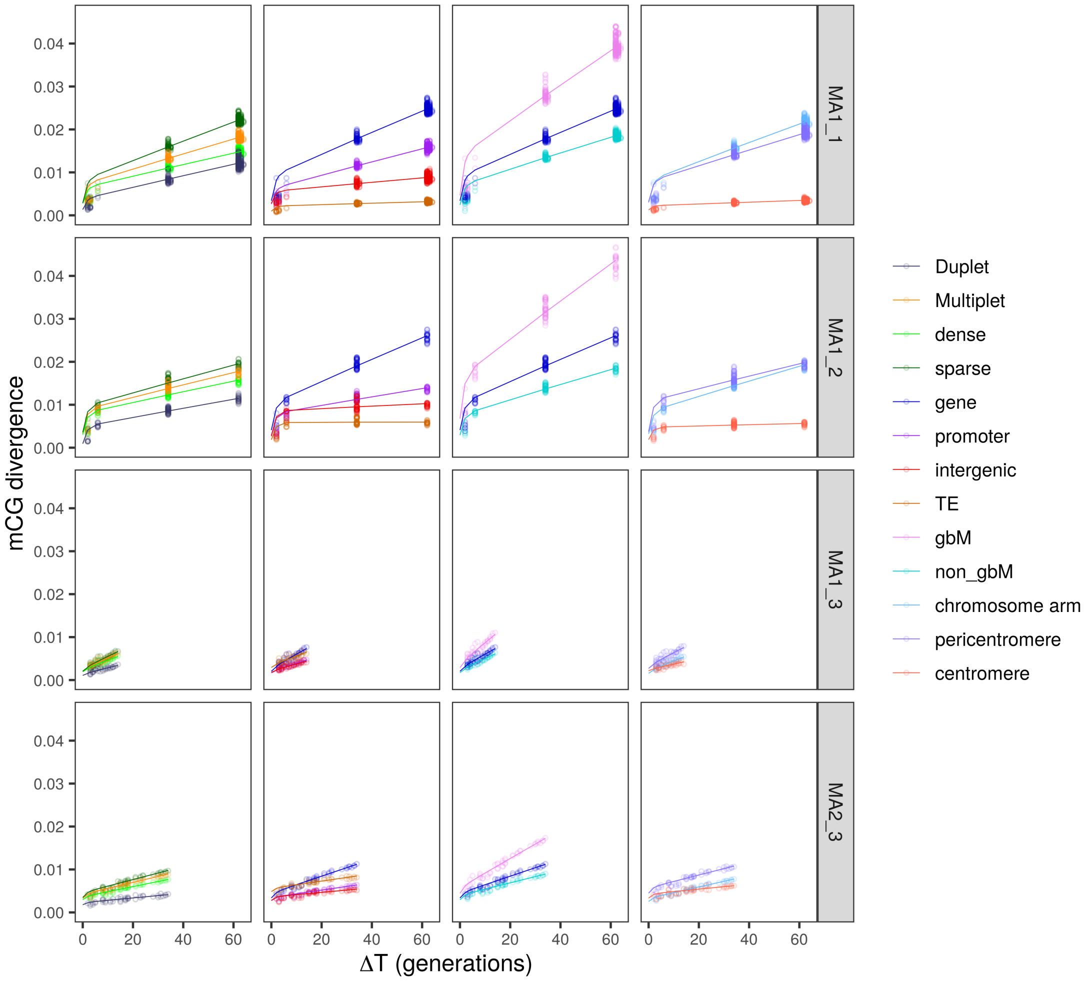


**Figure S4:** Divergence for all investigated genomic features and MA line pedigrees.

| | type | MA1_1.rates | MA1_2.rates | MA1_3.rates | MA2_3.rates | MA1_1.SE | MA1_2.SE | MA1_3.SE | MA2_3.SE |
|---|---|---|---|---|---|---|---|---|---|
| genome-wide | | $1.29\cdot10^{-4}$ | $1.06\cdot10^{-4}$ | $1.66\cdot10^{-4}$ | $7.84\cdot10^{-5}$ | $2.35\cdot10^{-6}$ | $4.10\cdot10^{-6}$ | $4.26\cdot10^{-5}$ | $2.44\cdot10^{-6}$ |
| gene | | $1.58\cdot10^{-4}$ | $1.65\cdot10^{-4}$ | $1.94\cdot10^{-4}$ | $1.14\cdot10^{-4}$ | $2.66\cdot10^{-6}$ | $4.14\cdot10^{-6}$ | $5.21\cdot10^{-5}$ | $3.27\cdot10^{-6}$ |
| gbM | | $3.09\cdot10^{-4}$ | $3.52\cdot10^{-4}$ | $3.38\cdot10^{-4}$ | $2.30\cdot10^{-4}$ | $5.08\cdot10^{-6}$ | $9.43\cdot10^{-6}$ | $4.62\cdot10^{-5}$ | $8.22\cdot10^{-6}$ |
| non-gbM | | $1.12\cdot10^{-4}$ | $1.06\cdot10^{-4}$ | $1.50\cdot10^{-4}$ | $8.05\cdot10^{-5}$ | $2.02\cdot10^{-6}$ | $3.04\cdot10^{-6}$ | $1.71\cdot10^{-5}$ | $2.96\cdot10^{-6}$ |
| promoter | | $9.36\cdot10^{-5}$ | $6.06\cdot10^{-5}$ | $1.04\cdot10^{-4}$ | $4.71\cdot10^{-5}$ | $1.87\cdot10^{-6}$ | $3.01\cdot10^{-6}$ | $2.47\cdot10^{-5}$ | $2.25\cdot10^{-6}$ |
| intergenic | | $5.64\cdot10^{-5}$ | $2.09\cdot10^{-5}$ | $9.79\cdot10^{-5}$ | $3.10\cdot10^{-5}$ | $3.44\cdot10^{-6}$ | $4.95\cdot10^{-6}$ | $9.45\cdot10^{-6}$ | $2.84\cdot10^{-6}$ |
| TE | $\alpha$ | $4.51\cdot10^{-5}$ | $1.28\cdot10^{-6}$ | $2.57\cdot10^{-4}$ | $9.60\cdot10^{-5}$ | $2.66\cdot10^{-6}$ | $8.73\cdot10^{-6}$ | $2.84\cdot10^{-5}$ | $8.14\cdot10^{-6}$ |
| 5'UTR | | $7.67\cdot10^{-6}$ | $1.07\cdot10^{-5}$ | $2.06\cdot10^{-5}$ | $1.55\cdot10^{-5}$ | $9.40\cdot10^{-7}$ | $9.60\cdot10^{-7}$ | $7.28\cdot10^{-6}$ | $1.25\cdot10^{-6}$ |
| 3'UTR | | $1.09\cdot10^{-4}$ | $1.06\cdot10^{-4}$ | $1.74\cdot10^{-4}$ | $6.54\cdot10^{-5}$ | $2.96\cdot10^{-6}$ | $2.90\cdot10^{-6}$ | $8.63\cdot10^{-5}$ | $4.02\cdot10^{-6}$ |
| chromosome arm | | $1.35\cdot10^{-4}$ | $1.09\cdot10^{-4}$ | $1.35\cdot10^{-4}$ | $7.30\cdot10^{-5}$ | $1.93\cdot10^{-6}$ | $3.01\cdot10^{-6}$ | $4.04\cdot10^{-5}$ | $2.28\cdot10^{-6}$ |
| pericentromere | | $1.44\cdot10^{-4}$ | $1.18\cdot10^{-4}$ | $1.97\cdot10^{-4}$ | $9.41\cdot10^{-5}$ | $2.71\cdot10^{-6}$ | $5.81\cdot10^{-6}$ | $3.06\cdot10^{-5}$ | $4.68\cdot10^{-6}$ |
| centromere | | $5.66\cdot10^{-5}$ | $4.03\cdot10^{-5}$ | $2.03\cdot10^{-4}$ | $8.87\cdot10^{-5}$ | $3.88\cdot10^{-6}$ | $8.67\cdot10^{-6}$ | $3.24\cdot10^{-5}$ | $9.43\cdot10^{-6}$ |
| genome-wide | | $2.83\cdot10^{-4}$ | $2.30\cdot10^{-4}$ | $8.74\cdot10^{-4}$ | $4.40\cdot10^{-4}$ | $5.16\cdot10^{-6}$ | $8.88\cdot10^{-6}$ | $2.23\cdot10^{-4}$ | $1.37\cdot10^{-5}$ |
| gene | | $9.50\cdot10^{-4}$ | $7.89\cdot10^{-4}$ | $1.48\cdot10^{-3}$ | $9.28\cdot10^{-4}$ | $1.60\cdot10^{-5}$ | $1.99\cdot10^{-5}$ | $4.03\cdot10^{-4}$ | $2.66\cdot10^{-5}$ |
| gbM | | $7.36\cdot10^{-4}$ | $6.95\cdot10^{-4}$ | $9.31\cdot10^{-4}$ | $6.89\cdot10^{-4}$ | $1.21\cdot10^{-5}$ | $1.87\cdot10^{-5}$ | $1.28\cdot10^{-4}$ | $2.47\cdot10^{-5}$ |
| non-gbM | | $1.27\cdot10^{-3}$ | $9.30\cdot10^{-4}$ | $2.43\cdot10^{-3}$ | $1.33\cdot10^{-3}$ | $2.30\cdot10^{-5}$ | $2.68\cdot10^{-5}$ | $2.81\cdot10^{-4}$ | $4.93\cdot10^{-5}$ |
| promoter | | $5.92\cdot10^{-4}$ | $3.16\cdot10^{-4}$ | $1.24\cdot10^{-3}$ | $5.88\cdot10^{-4}$ | $1.19\cdot10^{-5}$ | $1.57\cdot10^{-5}$ | $2.98\cdot10^{-4}$ | $2.82\cdot10^{-5}$ |
| intergenic | | $4.86\cdot10^{-5}$ | $4.05\cdot10^{-5}$ | $5.66\cdot10^{-4}$ | $1.92\cdot10^{-4}$ | $2.96\cdot10^{-6}$ | $9.60\cdot10^{-6}$ | $5.48\cdot10^{-5}$ | $1.76\cdot10^{-5}$ |
| TE | $\beta$ | $1.01\cdot10^{-5}$ | $3.20\cdot10^{-7}$ | $1.95\cdot10^{-4}$ | $7.63\cdot10^{-5}$ | $6.00\cdot10^{-7}$ | $2.19\cdot10^{-6}$ | $2.15\cdot10^{-5}$ | $6.48\cdot10^{-6}$ |
| 5'UTR | | $4.51\cdot10^{-4}$ | $4.82\cdot10^{-4}$ | $1.65\cdot10^{-3}$ | $1.29\cdot10^{-3}$ | $5.56\cdot10^{-5}$ | $4.32\cdot10^{-5}$ | $5.91\cdot10^{-4}$ | $1.04\cdot10^{-4}$ |
| 3'UTR | | $1.56\cdot10^{-3}$ | $1.37\cdot10^{-3}$ | $4.73\cdot10^{-3}$ | $1.86\cdot10^{-3}$ | $4.24\cdot10^{-5}$ | $3.76\cdot10^{-5}$ | $2.43\cdot10^{-3}$ | $1.16\cdot10^{-4}$ |
| chromosome arm | | $8.86\cdot10^{-4}$ | $6.43\cdot10^{-4}$ | $1.51\cdot10^{-3}$ | $8.75\cdot10^{-4}$ | $1.26\cdot10^{-5}$ | $1.79\cdot10^{-5}$ | $4.58\cdot10^{-4}$ | $2.74\cdot10^{-5}$ |
| pericenrtomere | | $2.62\cdot10^{-4}$ | $1.99\cdot10^{-4}$ | $7.82\cdot10^{-4}$ | $3.92\cdot10^{-4}$ | $4.94\cdot10^{-6}$ | $9.77\cdot10^{-6}$ | $1.22\cdot10^{-4}$ | $1.95\cdot10^{-5}$ |
| centromere | | $1.25\cdot10^{-5}$ | $8.32\cdot10^{-6}$ | $9.41\cdot10^{-5}$ | $4.25\cdot10^{-5}$ | $8.60\cdot10^{-7}$ | $1.79\cdot10^{-6}$ | $1.50\cdot10^{-5}$ | $4.52\cdot10^{-6}$ |
| genome-wide | | 2.19 | 2.18 | 5.28 | 5.61 | 0 | 0 | 0 | 0 |
| gene | | 5.99 | 4.79 | 7.63 | 8.1 | 0 | 0 | 0 | 0 |
| gbM | | 2.38 | 1.98 | 2.75 | 2.99 | 0 | 0 | 0 | 0 |
| non-gbM | | 11.32 | 8.78 | 16.21 | 16.55 | 0 | 0 | 0 | 0 |
| promoter | | 6.32 | 5.21 | 12 | 12.48 | 0 | 0 | 0 | 0 |
| intergenic | | 0.86 | 1.94 | 5.78 | 6.19 | 0 | 0 | 0 | 0 |
| TE | $\beta/\alpha$ | 0.22 | 0.25 | 0.76 | 0.79 | 0 | 0 | 0 | 0 |
| 5'UTR | | 58.83 | 44.95 | 80.11 | 82.95 | 0 | 0 | 0 | 0 |
| 3'UTR | | 14.24 | 12.9 | 27.18 | 28.49 | 0 | 0 | 0 | 0 |
| chromosome arm | | 6.54 | 5.92 | 11.22 | 11.99 | 0 | 0 | 0 | 0 |
| pericenrtomere | | 1.82 | 1.68 | 3.97 | 4.17 | 0 | 0 | 0 | 0 |
| centromere | | 0.22 | 0.21 | 0.46 | 0.48 | 0 | 0 | 0 | 0 |

**Table S1:** Epimutation rates estimates for regions genome-wide and regions overlapping with different annotations. mCG gain rate ($\alpha$), mCG loss rate ($\beta$) and $\beta/\alpha$ ratio per MA line. The last 4 column detail the standard error (SE) associated with each rate. These are calculated from bootstrapping.
